# Transcriptional Characterization of Nuclear-Integrated Organellar DNA in *Populus*

**DOI:** 10.64898/2026.07.08.737317

**Authors:** Reed Arneson, William Wittstock, Aimee Marceau, Yinan Yuan

## Abstract

The continuous transfer of organellar DNA into the nuclear genome during eukaryotic evolution has resulted in the widespread occurrence of nuclear plastid DNA insertions (NUPTs) and nuclear mitochondrial DNA insertions (NUMTs). However, their functional significance in nuclear gene expression and genome evolution remains largely unresolved. In this study, we employed Oxford Nanopore Direct RNA Sequencing (DRS) to investigate the transcription of NUPTs and NUMTs in the *Populus* nuclear genome and compared their transcriptional characteristics with their genome-wide insertion patterns. Our analyses revealed that the majority of transcribed NUPTs and NUMTs are enriched within introns and are co-transcribed with their host or adjacent genes in polycistronic-like transcriptional units. In addition, NUPTs and NUMTs frequently generate intronless transcripts, features reminiscent of their prokaryotic ancestry. We further identified a putatively functional NUPT-derived *psbH* gene that is unique to *P. trichocarpa*, providing new insights into the evolution of nuclear-encoded organelle-targeted genes. In addition, we identified transcribed NUPT and NUMT insertion polymorphisms among alleles, suggesting that organellar DNA insertions contribute to allelic variation and may participate in environmental adaptation. Collectively, our findings reveal previously unrecognized roles of NUPT and NUMT transcription in gene regulation, allelic variation, genome evolution, and the emergence of novel genes.

## INTRODUCTION

In eukaryotes, endosymbiotic gene transfer (EGT) has resulted in the relocation of numerous organellar genes to the nuclear genome [1]. This process is believed to be ongoing, with organellar DNA fragments continuously transferred into the nuclear genome [2-4]. Most of these transferred fragments are thought to be noncoding sequences and are retained as nuclear integrants of plastid DNA (NUPTs) and nuclear integrants of mitochondrial DNA (NUMTs) [5]. The persistence of organellar DNA transfer long after the establishment of endosymbiosis suggests that NUPTs and NUMTs may continue to serve unresolved biological functions. The widespread integration of organelle-derived DNA in many eukaryotic genomes is thought to be a major driver of nuclear genome evolution by providing raw genetic material for the emergence of novel genes and regulatory elements [6, 7]. However, the molecular mechanisms by which NUPTs and NUMTs contribute to genome evolution remain largely unknown.

Large numbers of NUPTs and NUMTs have been identified in many plant genomes [8, 9]. While numerous studies have characterized their genomic distribution, insertion patterns, and evolutionary dynamics across species, investigations into their transcription remain scarce and inconclusive. Consequently, NUPTs and NUMTs are generally regarded as transcriptionally inactive [10, 11], and their potential functional roles have largely been overlooked. One major technical challenge in studying the transcription of nuclear-integrated organellar DNA is its high sequence similarity to the corresponding organellar genomes [11]. With conventional short-read RNA sequencing, it is often difficult to distinguish whether transcripts originate from nuclear-integrated organellar sequences or from the chloroplast or mitochondrial genomes themselves. As a result, transcripts derived from organellar genomes may be incorrectly assigned to NUPT or NUMT loci in the nuclear genome, leading to ambiguous or misleading conclusions.

In this study, we employed Oxford Nanopore Direct RNA Sequencing (DRS), which directly sequences native RNA molecules at the transcript level [12], in *P. trichocarpa* (Nisqually-1). The long-read DRS data enabled the accurate identification of nuclear-integrated organellar DNA-derived transcripts that uniquely map to the nuclear genome despite their high sequence homology with organellar genomes. Leveraging this technological advantage, we characterized the transcriptional landscape of NUPTs and NUMTs in the *Populus* nuclear genome, including their expression patterns, allelic variation, and the identification of novel transcriptional units.

## RESULTS

### Insertions of NUPTs and NUMTs in *Populus* Genome

Organellar DNA insertions in the *Populus* nuclear genome were identified using RepeatMasker, a homology-based approach that uses organellar genome sequences as the libraries [7]. For NUPT identification, we used the *P. trichocarpa* chloroplast genome as the library (METHODs). Because a complete *P. trichocarpa* mitochondrial genome sequence is not yet published, we used the mitochondrial genome of hybrid poplar 717 as the reference library for NUMT identification. Given the limited synteny (>97%) of mitochondrial genomes among *Populus* species [13], despite they share high sequence homology (>96% identity), we interpreted the NUMT results with appropriate caution.

We identified 3,587 plastid DNA insertions (NUPTs), totaling 824,816 bp and accounting for approximately 0.21% of the *P. trichocarpa* nuclear genome, and 9,690 mitochondrial DNA insertions (NUMTs), totaling 2,558,388 bp and accounting for approximately 0.65% of the nuclear genome. This distribution is consistent with previous reports in other species showing that NUMTs are more abundant than NUPTs in nuclear genomes [8]. The median insertion lengths were 115 nucleotides (nt) for NUPTs and 150 nt for NUMTs (Fig. 1A). The largest NUPT was a 48,782-bp insertion located on chromosome 13, whereas the largest identified NUMT was a 62,127-bp insertion on chromosome 7. Because NUMTs were identified using the 717 mitochondrial genome as the library, the size of the largest NUMT is likely underestimated, and larger NUMT insertions may be present in the *P. trichocarpa* genome. Overall, 75% of NUPTs and NUMTs shared less than 90% sequence similarity with their organellar genome counterparts. In contrast, the remaining 25% shared more than 90% sequence similarity with their organellar counterparts. Notably, portions of the larger insertion blocks exhibited over 99% sequence identity to their corresponding organellar genome sequences, indicating that these insertions likely represent relatively recent transfer events, consistent with observations in other plant species.

**Figure 1.**
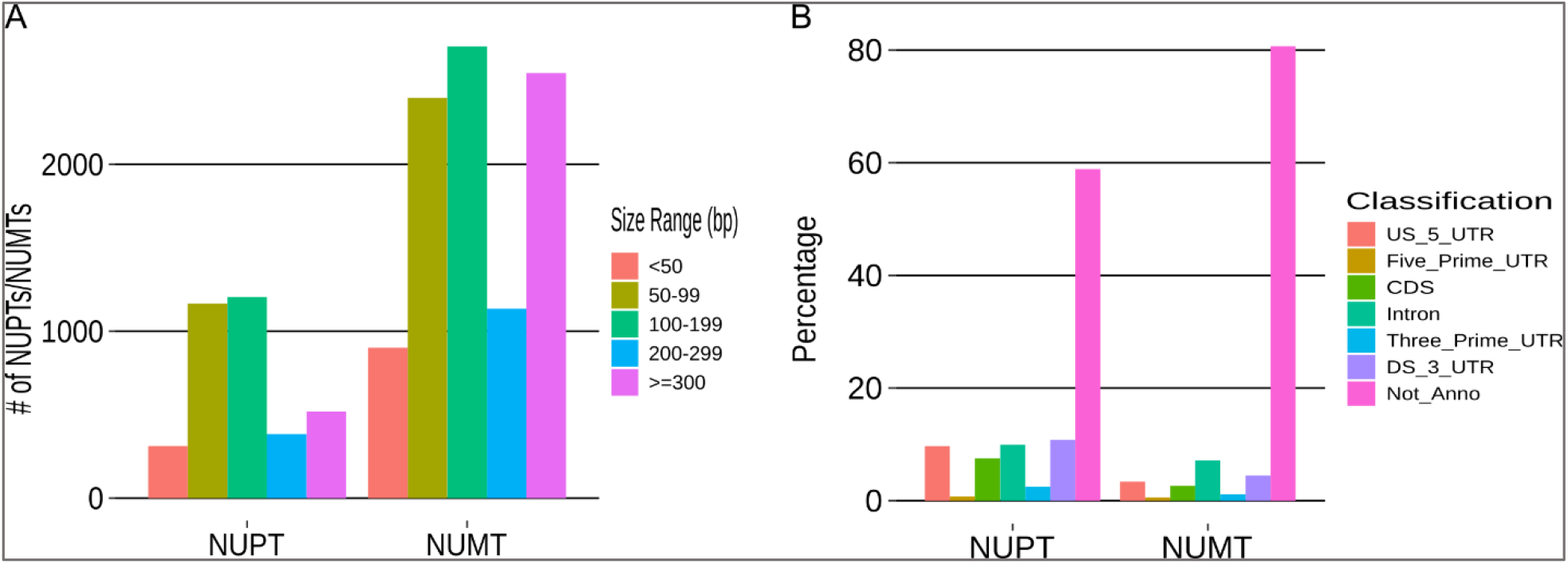
Size range and genome-wide distribution of NUPT and NUMT insertions in the *P. trichocarpa* nuclear genome. **A**. Size range of nuclear inserted organellar DNAs. **B**. Distribution of NUPTs and NUMTs across genomics regions. Bars represent the percentage of NUPTs and NUMTs located in different genomic features, including the 1-kb upstream region of 5′ UTR of annotated genes (US_5_UTR), 5′ UTR (Five_Prime_UTR), coding sequences (CDS), introns, 3′ UTR (Three_Prime_UTR), the 1 kb downstream of 3′ UTR of annotated genes (DS_3_UTR), and intergenic regions or not annotated genomic regions (Not_Anno). Percentages are calculated relative to the total number of genome-wide NUPT or NUMT insertions, respectively.

We next examined the genomic distribution of NUPTs and NUMTs (Fig. 1B). More than 60% of NUPTs and over 80% of NUMTs were located in unannotated intergenic regions. Among insertions within annotated genes, NUMTs were most frequently found in introns, followed by the 1-kb upstream regions of the 5′ untranslated regions (5′ UTRs) and the 1-kb downstream regions of the 3′ untranslated regions (3′ UTRs), and were least frequent within the 5′ and 3′ UTRs themselves. In contrast, NUPTs were distributed more evenly among introns, the 1-kb upstream regions of 5′ UTRs, and the 1-kb downstream regions of 3′ UTRs.

### Distinct Transcriptional Features of NUPTs and NUMTs

The distribution of transcribed NUPTs and NUMTs across genomic regions differed markedly from their genomic insertion patterns. Although our previous analysis showed that most NUPTs and NUMTs are inserted into unannotated intergenic regions, transcribed NUPTs and NUMTs, particularly NUPTs, were strongly associated with annotated genes. Most transcribed NUPTs were co-transcribed with their host or neighboring genes on the same RNA molecules (Fig. 2A-D). Among the 111 transcribed NUPTs identified, approximately 67% were located within introns of annotated genes and were incorporated into intron-retained transcripts (Fig. 2A, 2E). An additional 14% were located within 3′ UTRs (Fig. 2B, 2E), whereas 12% were located downstream of annotated 3′ UTRs (Fig. 2C, 2E) but were transcriptionally linked to neighboring genes on same RNA molecules. Smaller proportions of transcribed NUPTs were detected within exons, 5′ UTRs, and the 1-kb upstream regions of annotated 5′ UTRs linking with the neighboring genes (Fig. 2D).

**Figure 2.**
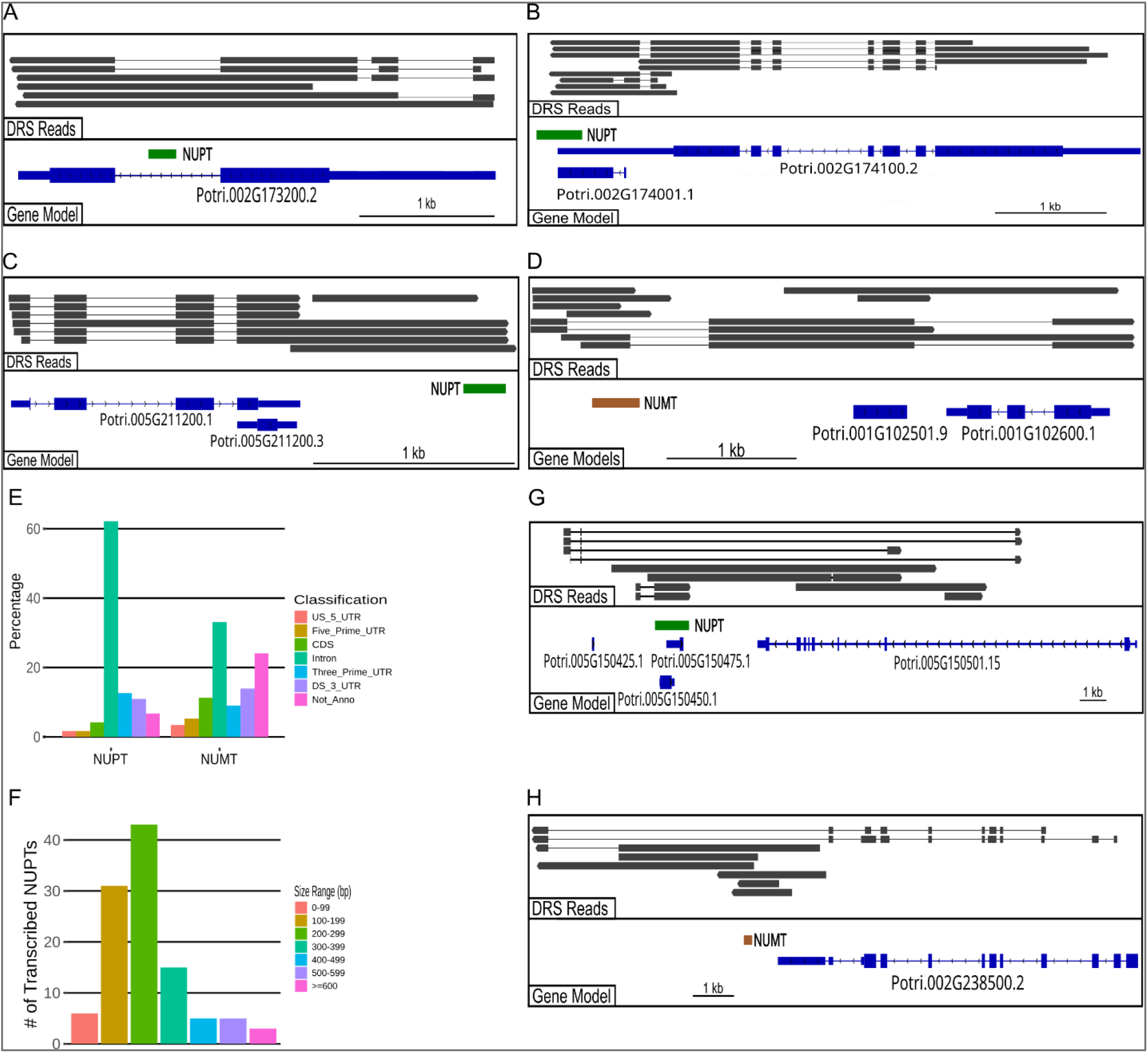
Transcriptional characteristics of NUPTs and NUMTs. (**A–D**) Representative examples of NUPT and NUMT co-transcription with annotated genes. **A**. Co-transcription through intron retention. **B**. Transcription in 3′ UTR. **C**. Transcription extending beyond the annotated 3′ UTR into the downstream region. **D**. Transcription through upstream of the annotated 5′ UTR. **E**. Transcribed NUPTs and NUMTs spanning across annotated genomic features. **F**. Size distribution of transcribed NUPT fragments. **G**. Long intronless NUPT transcripts are detected in a predicted intron of unannotated genomic region. **H**. Intronless NUPT transcripts detected downstream of 3′ UTR region of one annotated gene.

The transcriptional patterns of NUMTs were more diverse than those of NUPTs. As shown in Fig. 2E, among the 254 transcribed NUMTs identified, approximately 25% were not associated with genes, whereas the remaining 75% of NUMTs, associated with annotated genes on the same RNA molecules.

As shown in Fig. 2E, the major co-transcriptional events included intron retention (35% of transcribed NUMTs) and transcription extending beyond the 3′ UTR (15% of transcribed NUMTs). Within annotated gene regions, NUMT transcription was most frequently detected in 3′ UTRs (9%), compared with exons, 5′ UTRs, and the 1-kb upstream regions of annotated 5′ UTRs. Overall, transcribed NUPTs and NUMTs shared 70-99% sequence identity with their respective organellar genome counterparts, with a median transcript length of 221.5 nt for NUPTs (Fig. 2F). Interestingly, the organellar DNA sequences represented in NUPTs were derived from both genic and intergenic regions of the chloroplast genome, whereas those represented in NUMTs originated predominantly from intergenic regions of the mitochondrial genome. Notably, nearly half of the transcribed NUPTs were associated with annotated genes involved in membrane or organelle functions.

A notable finding from the transcription analysis is that NUPT and NUMT transcripts are frequently incorporated into transcripts of their host or neighboring genes. In both cases, we found that the transcribed NUPTs and NUMTs contribute minimally to the exonic regions of annotated genes (Fig. 2E). However, their strong enrichment in intron retention, particularly among transcribed NUPTs, is noteworthy. Intron retention is a major form of alternative splicing. Once considered merely transcriptional noise, it is now recognized as a sophisticated post-transcriptional regulatory mechanism that contributes to gene regulation, transcriptome diversification, and rapid cellular responses. Given the enrichment of NUPTs and NUMTs in retained introns, the functional significance of these intron retention events remains to be investigated.

Additionally, the co-transcription of NUPTs and NUMTs with neighboring genes is noteworthy. When NUPTs or NUMTs are located adjacent to annotated genes, they are frequently co-transcribed with their neighboring genes, generating transcript structures reminiscent of polycistronic transcription (Fig. 2C-D). Notably, many of these NUPTs and NUMTs are also transcribed independently, producing both monocistronic transcripts and polycistronic-like RNAs.

As noted above, the majority of transcribed NUPTs and NUMTs associated with host genes were incorporated into host gene transcripts through intron retention. In addition, we observed that nuclear-inserted organellar DNA sequences frequently generated intronless transcripts regardless of their genomic location, including both coding and non-coding regions (Fig. 2G-H). This transcriptional pattern was particularly common among transcribed NUMTs located in unannotated genomic regions. The occurrence of polycistronic-like transcripts together with the frequent intronless transcription is reminiscent of the prokaryotic mode of gene expression, suggesting that NUPTs and NUMTs may retain transcriptional characteristics inherited from their prokaryotic ancestors.

In our study, transcripts containing the organellar DNA sequence are polyadenylated [poly(A)] and most possess poly(A) tails longer than 60 nucleotides, suggesting they are likely to be integrated with nuclear gene expression and unlikely to be destined for nonsense-mediated decay.

### Characterization of a Transcribed *psbH*-Derived NUPT

We also reidentified a previously reported polycistronic-like NUPT from our group that shares 94% nucleotide sequence identity with the chloroplast *psbH* gene [14]. *psbH* encodes an essential component of Photosystem II and, to our knowledge, is exclusively encoded by the chloroplast genome in all known plant species. Unlike most NUPTs, which are non-coding organellar DNA fragments, this nuclear *psbH* copy contains a predicted intact open reading frame (Fig. 3A). Sequence analysis also identified a putative plastid-targeting transit peptide at its 5′ end, indicating that the encoded protein is likely targeted to the chloroplast. Thus, *P. trichocarpa* contains two *psbH* genes: one encoded by the chloroplast genome and one by the nuclear genome. Notably, we previously reported that the nuclear *psbH* gene is predominantly co-transcribed with its host gene on the same RNA molecules [14], suggesting that it may have acquired a function distinct from that of the chloroplast-encoded *psbH* gene.

**Figure 3.**
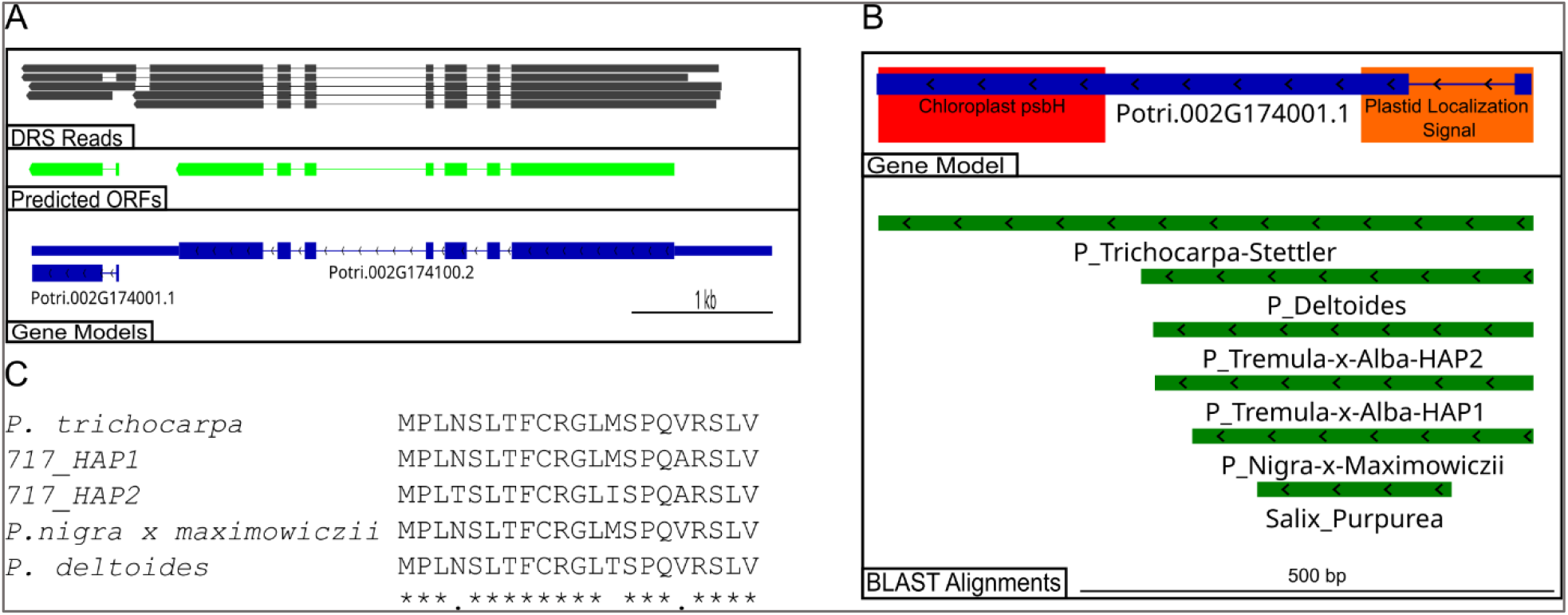
Characterization of a novel NUPT encoding a nuclear *psbH* gene. **A**. Gene structure and predicted open reading frame (ORF) of the nuclear *psbH* gene. **B**. Comparative analysis of the genomic region containing the nuclear *psbH* locus across selected *Populus* species. **C**. Conservation of the plastid-targeting transit peptide amino acid sequence among the *Populus* genomes examined.

To examine the distribution of this nuclear *psbH* copy, we searched additional *Populus* genomes with available genome assemblies. We focused on the 13.5 kb genomic region containing the *psbH* NUPT, its host gene Potri.002G174100, which encodes a WD40 repeat protein, and the downstream gene Potri.002G173900, which encodes a putative MYB transcription factor. Across all *Populus* genomes examined, the two flanking genes (WD40 and MYB) were highly conserved, whereas the intervening region containing the *psbH* NUPT exhibited substantial sequence variation. The sequence corresponding to chloroplast *psbH* was detected only in *P. trichocarpa* genome, including the ‘Stettler-14’ genotype, and was absent from all other *Populus* genomes and one *Salix* (willow) genome examined (Fig. 3B). In contrast, the sequence encoding the predicted plastid-targeting transit peptide (21 amino acids; Fig. 3C) was conserved at the corresponding genomic region in all *Populus* genomes surveyed but was absent the willow genome. Although the corresponding genomic region in willow shares approximately 89% sequence identity with *Populus*, it lacks the first 30 nucleotides of this region and therefore does not encode a plastid-targeting transit peptide (Fig. 3B). These findings suggest a complex evolutionary relationship between the acquisition of the organelle-targeting peptide sequence and organelle-to-nucleus gene transfer in *Populus*.

### NUPTs and NUMTs as a Source of Allelic Variation

While examining the transcriptional patterns of NUPTs and NUMTs, we found that some transcribed organellar DNA sequences did not align to the *P. trichocarpa* reference genome, whereas the flanking non-organellar sequences within the same RNA molecules aligned. We hypothesized that these NUPT/NUMT sequences originated from alleles that were not represented in the current *P. trichocarpa* reference genome assembly. To test this hypothesis, we designed sequence-specific primers flanking two NUPT insertion sites for genomic DNA amplification. As expected, PCR amplification from Nisqually-1 genomic DNA yielded the predicted NUPT-containing products, together with shorter amplicons consistent with alleles lacking the corresponding NUPT/NUMT insertion (Fig. 4). The identities of the NUPT/NUMT-containing amplicons were confirmed by cloning and Sanger sequencing. These results confirm that the allelic differences at the two annotated genomic loci examined were attributable to the presence or absence of NUPT insertions. In both loci, the NUPT insertion was located within an intron.

**Figure 4.**
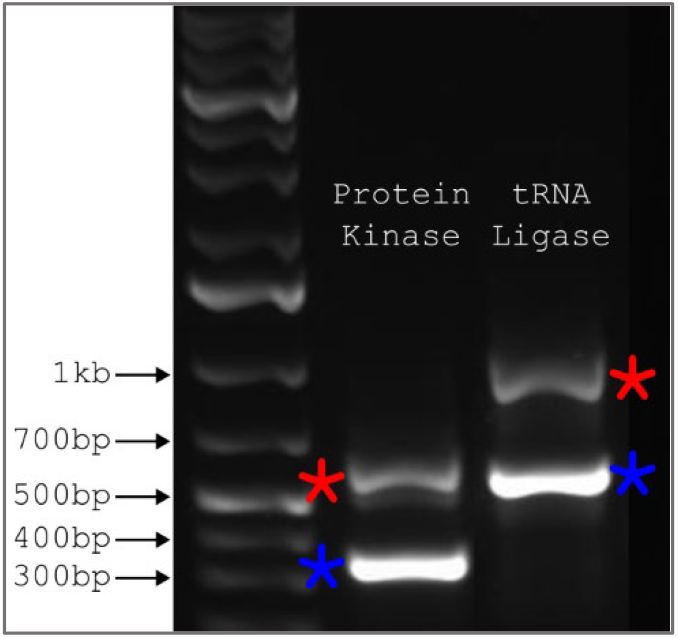
PCR validation of NUPT-containing and NUPT-lacking alleles of Potri.003G049700 (protein kinase) and Potri.007G129800 (tRNA ligase). PCR amplicons correspond to the allele containing the NUPT insertion (red asterisk) or lacking the insertion (blue asterisk) are indicated.

To note, besides the presence or absence of NUPT and NUMT insertions, we also identified single-nucleotide polymorphisms (SNPs) and short deletions within NUPT and NUMT sequences as common forms of allelic variation. In addition, allelic sequence variation was frequently observed in genomic regions flanking NUPT and NUMT insertions.

Overall, our results demonstrate that the insertion of NUPTs and NUMTs can contribute to allelic variation and influence allele-specific transcription, providing new insights into the potential functional significance of the widespread occurrence of NUPTs and NUMTs in eukaryotic nuclear genomes and complementing existing theories regarding their role in the evolution of novel genes.

## DISCUSSION

Throughout eukaryotic evolution, fragments of organellar genomes have been continuously transferred to the nuclear genome through mechanisms that remain poorly understood. The functional significance of these nuclear-integrated organellar DNA sequences also remains largely unresolved. Numerous studies have characterized the genomic distribution of NUPTs and NUMTs in an effort to understand their biological roles. These studies have shown that large NUPT and NUMT insertions are frequently enriched in centromeric regions, whereas the genomic distribution of smaller insertions varies among species [8, 9].

NUPTs and NUMTs have generally been regarded as transcriptionally inactive components of the nuclear genome [15]. Although they represent a rich source of genetic material for genome evolution, the mechanisms by which they contribute to this process remain poorly understood. Genomic analyses have provided valuable insights into the distribution and evolutionary history of NUPTs and NUMTs but have offered only limited information about their functional roles. In this study, we adopted a complementary approach by integrating transcriptional analyses with genomic characterization to investigate the expression patterns of NUPTs and NUMTs and to assess their potential contributions to gene regulation, allelic variation, and the evolution of novel genes.

### Impacts of NUPTs and NUMTs on Gene Regulation and the Evolution of Novel Transcriptional Units and Alleles

Our findings provide new insights into the potential impact of organellar DNA transfer on the evolution of gene regulation and transcriptional innovation. First, transcribed NUPTs and NUMTs were highly enriched within introns, and the majority were co-transcribed with their host genes as part of intron retention events. Given that intron retention is an important mechanism of post-transcriptional gene regulation [16, 17], these observations suggest that the transcription of NUPTs and NUMTs may influence host gene regulation through intron retention.

Second, the polycistronic-like transcription of NUPTs and NUMTs observed in this study reveals a previously unrecognized transcriptional association between organellar DNA insertions and adjacent nuclear genes. Whether this co-transcription influences the expression of host or neighboring genes, or instead reflects transcription initiated from those genes, remains to be determined. Regardless of the underlying mechanism, however, these events generate novel polycistronic-like transcriptional units in the *Populus* nuclear genome, thereby expanding its transcriptional repertoire.

Another notable finding is that NUPTs and NUMTs are present in different alleles at some genomic loci, thereby contributing to allelic variation. Given the high levels of heterozygosity and frequent interspecific introgression in natural *Populus* populations [18], insertion polymorphisms of NUPTs and NUMTs may represent an additional source of allelic diversity that is introduced and maintained through hybridization and introgression. Such allelic variants may contribute to nuclear genome evolution by originating through organellar DNA transfer, becoming fixed within nuclear genomes, and subsequently spreading among populations or species via hybridization and introgression, where they are subject to natural selection.

### Evolution of Nuclear-Encoded Organelle-Targeted Genes

Throughout the coevolution of organelles and the nucleus, successful EGT has generated numerous nuclear-encoded, organelle-targeted proteins that are imported back into organelles to perform their functions. The acquisition of organelle-targeting signals has therefore been regarded as a critical step in this evolutionary process. It is generally assumed that these targeting signals evolved after organellar genes were transferred to the nuclear genome [19]. However, our findings raise an alternative possibility.

We found that the nuclear *psbH* copy is present only in *P. trichocarpa*, whereas the sequence encoding its predicted plastid transit peptide is conserved at the corresponding genomic region across all *Populus* species examined (Fig. 3B). This observation suggests two possible evolutionary scenarios. Under the conventional model, the transit peptide evolved after organellar gene transfer, implying that the nuclear *psbH* gene was subsequently lost in the other *Populus* lineages. Alternatively, the transit peptide-encoding sequence may have evolved before organellar gene transfer. In this scenario, the nuclear *psbH* gene in *P. trichocarpa* would represent a relatively recent NUPT insertion that integrated into a genomic locus already containing a functional targeting sequence, thereby giving rise to a potentially functional nuclear gene. If this scenario is correct, the evolution of organelle-targeted genes may not always require the de novo acquisition of targeting signals following gene transfer. Instead, pre-existing targeting sequences in the nuclear genome could occasionally facilitate the functionalization of newly transferred organellar genes. Determining how frequently this mechanism has contributed to the evolution of organelle-targeted genes will require further investigation.

## Conclusion

Collectively, our findings suggest that NUPTs and NUMTs retain transcriptional characteristics reminiscent of their prokaryotic ancestry while contributing to the evolution of nuclear gene regulation. In *Populus*, NUPTs and NUMTs are predominantly co-transcribed with host genes as polycistronic-like transcriptional units and frequently generate intronless transcripts, both of which resemble features of prokaryotic gene expression. Together with our previous finding that nuclear genes associated with organelle functions are enriched within polycistronic-like transcriptional units [14], these observations support the hypothesis that polycistronic-like transcription represents an evolutionarily derived mode of nuclear gene expression in plants.

More broadly, our results indicate that organellar DNA transfer has contributed not only genetic material but also novel patterns of transcriptional organization and gene regulation to plant nuclear genomes. By generating new transcriptional units, influencing gene regulation, and creating allelic variation, NUPTs and NUMTs may have played a broader role in shaping the structural and functional evolution of plant nuclear genomes than has previously been recognized.

## METHODS

### Plant Materials

*P. trichocarpa* Nisqually-1 plants used in this study were propagated from stem cuttings kindly provided by Dr. Tuskan at Oak Ridge National Laboratory. Plants were grown in 3-gallon pots filled with a professional growing medium (Sun Gro) in a greenhouse and maintained at full field capacity (FC) for two months before drought treatment [20]. For the drought treatment, irrigation was withheld until soil water content decreased to 40% FC, and plants were maintained at this level for three weeks. Control plants were watered regularly to maintain full FC throughout the experiment. Pots were weighed daily to monitor soil water content. Upon completion of drought treatment, both mature leaf (LPI 7) and root tissues were flash-frozen in liquid nitrogen for total RNA extraction.

### Nanopore DRS Library Construction, Sequencing, Data Acquisition and Processing

Nanopore DRS libraries were constructed as before following manufacturer’s instructions [14]. The libraries were sequenced on PromethION RNA flow cells (Oxford Nanopore Technologies, FLO-PRO004RA) using a PromethION 2 Solo (P2 Solo) device connected to a GridION Mk1 as a compute resource and operated with standard MinKNOW software (version 24.11.8).

Raw pod5 files generated from sequencer were basecalled using the Dorado software (version 0.9.2+2634e9f) provided by Oxford Nanopore Technologies. The basecalling was performed with the model <u>rna004_130bps_sup@v5.1.0</u>, along with poly(A) tail length estimation using the command: dorado basecaller sup --estimate-poly-a.

### Nanopore DRS reads mapping and identification of NUPTs and NUMTs transcripts

Basecalled BAM files were converted to FASTQ using SAMtools fastq command: samtools fastq. The resulting RNA reads were independently aligned to the *P. trichocarpa* reference genome v4.1 (nuclear alignment) and to the organellar genomes (organellar alignments), including the *P. trichocarpa* chloroplast genome (GenBank accession EF489041.1) and the *P. tremula* × *P. alba* mitochondrial genome (GenBank accession KT429213.1), using minimap2 [21] with the parameters: -a -x splice -k14 -uf. The resulting BAM files were filtered with SAMtools [22] using the command: samtools view -h -F 2324 to exclude unmapped, secondary, and supplementary reads.

To identify transcripts derived from nuclear-integrated organellar DNA sequences, we first filtered the nuclear BAM files to retain only reads that also aligned to the organellar genomes. To distinguish transcripts originating from NUPTs and NUMTs from authentic organellar RNAs, we used pysam to identify reads that exhibited at least 100 nt of soft clipping at either end in the organellar genome alignments while exhibiting less than 50 nt of soft clipping at both ends in the corresponding nuclear genome alignments. Reads meeting these criteria were retained as candidate NUPT- or NUMT-derived transcripts for downstream analyses.

### Identification of Genome-wide Insertion Pattern of NUPTs and NUMTs

NUPT and NUMT insertions were identified using RepeatMasker [7]. RepeatMasker was ran with the parameters: -pa 10 -no_is -no_low -norna -lib Organelle.fa on the *P. trichocarpa* reference genome (v4). The Organelle.fa file was a custom library composed of the *P. trichocarpa* chloroplast genome and *P. tremula* x *P. alba* mitochondrion genome to identify NUPTs and NUMTs, respectively. NUPTs and NUMTs identified from RepeatMasker were converted into separate BED files and each file was merged so that overlapping regions would only be counted once. The bed files were then analyzed in R to determine the number of insertions and the size distribution of each insertion type. The resulting data was graphed using ggplot2 in R.

To classify NUPTs and NUMTs to specific genomic features, bedtools intersect was used to determine the number of bases in NUPTs and NUMTs overlapped with each genomic feature (5’ UTR, CDS, intron, 3’ UTR, 1 kb downstream region of an annotated gene, and intergenic regions). The graphs represent the percentage of bases for either NUPT or NUMTs associated with each feature. The data was graphed using ggplot2 in R.

### Genomic DNA extraction, PCR amplification, and Sanger sequencing

Genomic DNA was extracted from leaf tissue of *P. trichocarpa* Nisqually-1 using a 2% CTAB extraction buffer containing 2 M NaCl, 25 mM EDTA, 0.1 M Tris-HCl (pH 9.0), 2% PVP (K-30), and 2% 2-mercaptoethanol. The extracted DNA was treated with RNase A/T1 (Thermo Fisher Scientific, EN0551) to remove residual RNA and subsequently used as the template for PCR amplification of the NUPT insertion alleles.

PCR products were separated on a 1% TAE agarose gel, and the two amplified bands were excised and purified using a gel purification kit (NEB, T1120S). The purified fragments were cloned into a TOPO™ TA vector (Thermo Fisher Scientific, 450030) and subjected to Sanger sequencing.

### Data Access

Sequencing data generated in this study including raw FASTQ and BAM files have been submitted to NCBI Sequence Read Archive (SRA) under the BioProject accession number PRJNA1354876. All the custom codes used in this study are available at GitHub (https://github.com/Reedgarn).

## Acknowledgements

We would like to thank the Linux Group of Information Technology at Michigan Technological University for their assistance with software installation.

This work was supported by research seed grants from Michigan Technological University, the Ecological Science Center, and the College of Forest Resources and Environmental Science (start-up funds to Y.Y.) at Michigan Technological University, as well as the McIntire-Stennis Fund (USDA NIFA project #7008315 to Y.Y.).*Author contributions:* Y.Y.: conceptualization; Y.Y. and R.A.: methodology; R.A. and A.M.: data collection; Y.Y., R.A., and W.W.: data analysis; Y.Y., R.A., and W.W.: manuscript preparation.

